# Novel technology to develop non-transmissible live influenza virus vaccines through alterations in M2/M42 ion channel expression

**DOI:** 10.1101/2023.04.11.536210

**Authors:** Darrell R. Kapczynski, Karen Segovia, Klaudia Chrzastek, Carina Conceicao, David L. Suarez, Lonneke Vervelde, Paul Digard

**Author notes:** Corresponding authors: Darrell R. Kapczynski, Exotic & Emerging Avian Viral Diseases SEPRL, ARS, USDA, 934 College Station Rd Athens, Ga 30605. APHA Weybridge New Haw, Addlestone, Surrey UK KT15 3NB. Biochemistry Department 1st floor, Macewen Building Glasgow Royal Infirmary 84 Castle Street G4 0SF Glasgow (Carina Conceicao). Paul Digard Division of Infection and Immunity The Roslin Institute and Royal (Dick) School of Veterinary Studies University of Edinburgh Easter Bush, Midlothian, U.K. D.R.K., L.V., and P.D. contributed equally to this work.

## Abstract

Continued global outbreaks of H5Nx highly pathogenic avian influenza viruses (HPAIV) have been reported in poultry since the emergence of the Asian Goose/Guangdong lineage of HPAIV H5N1 in 1996. Consequently, vaccines have been developed and employed to protect commercial and non-commercial poultry flocks. However, constant changes in virus immunogenetics makes vaccine candidates that prevent morbidity, mortality, reduce shedding, and prevent transmission a moving target. Here, we tested the effects of disrupting the mRNA splice sites and thus expression of either of the two isoforms of the viral ion channel M2 and M42 expressed by the H5N2 low pathogenic avian influenza virus (LPAIV) progenitor of the 1983 Pennsylvania HPAIV outbreak. Both G52C (ΔM2) and G145A (ΔM42) versions of the A/chicken/Pennsylvania/1/83 (Ck/Penn) virus replicated well *in vitro* and *in ovo*, but showed altered virion morphology, with the G145A virus exhibiting filamentous budding. The G52C and G145A viruses also infected chickens efficiently and stimulated robust immune responses; however, unlike the wild type virus, they did not transmit to naïve-contact birds. In protection studies, vaccination of birds with G52C or G145A viruses protected 100% of birds from lethal challenge with homologous and distant heterologous H5 HPAIV strains from North American and Asian lineages and significantly reduced virus shedding compared to controls. Furthermore, the live virus vaccines decreased virus shedding 10,000-fold more than corresponding inactivated forms of the vaccine virus. Taken together, these studies demonstrate a strategy for developing a non-transmittable but highly protective live attenuated virus for AIV.

## Introduction

The genome of avian influenza virus (AIV) consists of eight RNA segments of negative polarity that encode 10 core proteins and a variable number of accessory polypeptides depending on virus strain ^1, 2^. Segment 7 encodes the viral matrix protein M1 as the primary gene product, as well as the M2 protein through mRNA splicing ^3^. M2, produced from mRNA2 (Figure 1A), is a 97 amino acid integral membrane protein with pH-gated ion channel activity that plays essential roles during the entry and egress of influenza A viral particles ^4^. Segment 7 from a laboratory-adapted human strain of influenza A virus (A/Wilson-Smith Neurotropic/33) has been shown to produce an additional spliced mRNA (mRNA4), initially predicted to encode an internally deleted version of M1 whose expression has not been detected ^5^. Subsequently, we showed that in another historical human strain of virus (A/PuertoRico/8/34; PR8), mRNA4 actually encoded, via leaky ribosomal scanning to access an internal AUG codon, an M2 isoform with a variant ectodomain (Figure 1B), denominated M42 ^6^. Subcellular localization analyses showed differences between M2 and M42 proteins, with M2 mainly localizing to the plasma membrane, while M42 was more perinuclear, concentrating around the Golgi apparatus ^6^. Most strains of influenza A virus are unlikely to produce M42, because the splice donor (SD) sequence needed to produce mRNA4 is not conserved; however, certain highly pathogenic avian influenza (HPAIV) strains, in particular the H5N2 virus (and its low pathogenic (LPAIV) progenitor) responsible for the major 1983 outbreak in Pennsylvania USA, represent an exception ^6, 7^. The biological significance of M42 versus M2 expression is unclear, but in the HPAIV background it was hypothesized that the Golgi-biased localization of M42 might aid in the intracellular transportation of the hemagglutinin (HA) protein, to prevent low pH-induced premature conformational change of the furin-cleavable HA ^6^.

**Fig 1.**
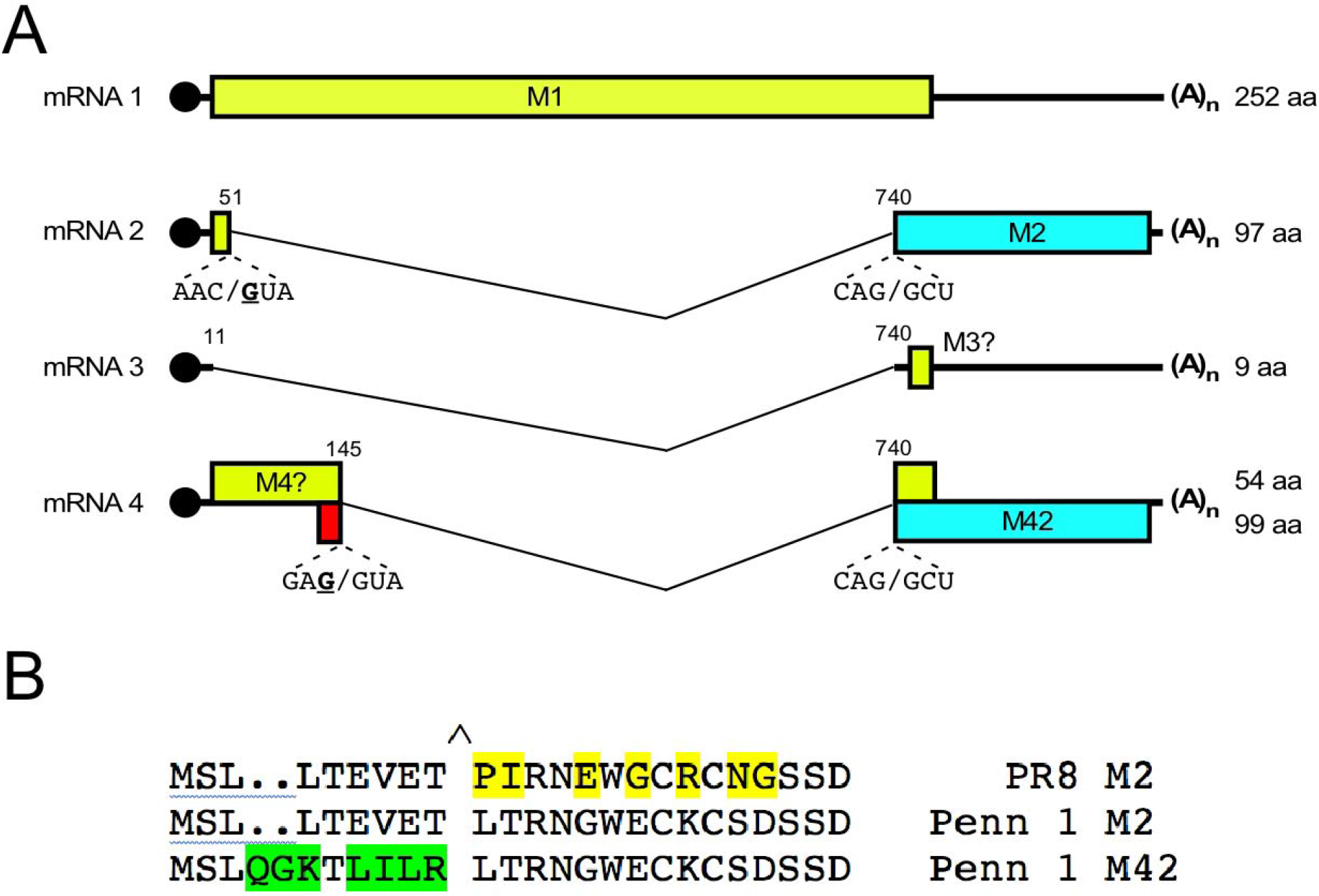
Segment 7 gene coding strategies. **(A)** The M segment has four transcripts: the non-spliced mRNA1 encodes the M1 protein starting from an open reading frame (ORF) running from nucleotides 26-28 until 782-784. Three alternatively spliced mRNAs are known, all using a common splice acceptor site at position 740, but 5’-splice donor sites at positions 11, (mRNA3), 51 (mRNA2) and 145 (mRNA4). M2 is translated from mRNA2, using the 5’-proximal AUG codon at positions 26-28 but then shifting reading frame at the splice site and terminating at a stop codon at nucleotides 1005-1007. M42 is produced from mRNA4 using an internal initiation codon at positions 114-116, changing reading frame at the splice site and terminating at the same stop codon as used for M2. The M3 and M4 ORFs in mRNAs 3 and 4 respectively encode hypothetical polypeptides derived from the M1 ORF. Colored boxes represent the coding regions, thin lines introns and thick lines the non-coding regions for each gene. Nucleotides mutated to disrupt splice donor sites are underlined in bold (Adapted from ^6^). (B) Ion channel ectodomain sequences of viruses used in this study. Residues varying from the Ck/Penn/1 M2 sequence are highlighted.

Infection with AIV can cause morbidity and high mortality in poultry and result in severe economic losses to poultry producers ^8, 9^. In 2014-2015 and again in 2021-2022, incursions of HPAI in the U.S. and Europe have resulted in devastating losses of poultry and wild birds in excess of 47 million per event due to infections and cullings ^9–11^. AIVs can also constitute a public health threat due to their zoonotic capacity ^12, 13^. Different measures have been adopted for preventing or controlling the disease in poultry. Territories free of HPAIV aim to eliminate the virus if infection occurs, through culling of affected birds and compensation schemes to poultry producers following an outbreak; whereas countries where HPAIV is endemic have installed vaccination protocols to protect susceptible animals and to reduce viral excretion and transmission following AIV challenge ^14–16^. Current poultry vaccination programs include the use of inactivated and recombinant subunit vaccines. Killed vaccines stimulate primarily a humoral immune response, in which anti-HA antibodies are protective ^17^. In contrast, live recombinant vaccines expressing the AIV HA and/or other proteins induce lower levels of humoral immunity, but can provide better protection through the induction of cellular immunity against the major viral structural proteins ^18, 19^. However, to achieve this, boosting with killed vaccine may be required ^14, 20, 21^. In addition, the breadth of protection of both these types of vaccine is restricted to the specific HA in the vaccine and they therefore do not protect against emerged or drifted strains ^22^. As a result, vaccinated birds still excrete detectable amounts of virus, display limited clinical signs of disease, and can serve as silent carriers ^23, 24^. Consequently, new technologies are being developed to increase the breadth of protection of AIV vaccines. One approach is the generation of live attenuated vaccines that can stimulate humoral, cellular and mucosal immunity against all viral proteins because of their ability to mimic the natural infection process ^25^. These types of vaccines are used for other poultry viral diseases, as well as human influenza; however, with concerns over genetic reassortment or transmission to off-target species.

Here, reverse genetics was utilized to generate viruses based on the LPAIV progenitor of the 1983 Pennsylvania outbreak to study the *in vitro* and *in vivo* role of the M2 and M42 proteins. We demonstrate that mutated viruses no longer expressing a mixture of M2 and M42 polypeptides showed significant alterations to virion size and morphology and although infectious *in vivo*, were unable to transmit to naïve birds. Furthermore, these non-transmissible viruses proved to be efficacious live-attenuated vaccines against homologous and heterologous HPAIV challenge. Taken together, this work demonstrates a role of M2/M42 altered expression on virus morphology and its impact on transmission in avian species, which can be exploited for the development of a live attenuated vaccine.

## Methods

### Ethics statement

All experimental bird procedures were approved and performed in strict accordance with the recommendation in the Guide for the Care and Use of Laboratory Animals and were reviewed and approved by the Southeast Poultry Research Laboratory (SEPRL) Institutional Animal Care and Use Committee (IACUC).

### Cells and viruses

DF-1 chicken fibroblast, HD-11 chicken macrophage-like, Madin-Darby Canine Kidney (MDCK) and 293T human embryonic kidney cells were cultured according to standard procedures ^26, 27^. Plasmid-based reverse genetics (rg) clones for the A/Puerto Rico/8/34 H1N1 (PR8) strain have been described ^28^. Similar plasmid clones of the A/chicken/Pennsylvania/1/1983 H5N2 LPAIV (Ck/Penn/WT) were generated using standard vectors ^28^. Briefly, seven Ck/Penn segments (PB1, PA, HA, NP, NA, M, and NS; GenBank accession numbers GU052777, GU052776, GU052771, GU052774, GU052773, GU052772, and GU052775 respectively) and the PB2 from A/Anhui/1/2013/H7N3 (GenBank accession number CY181528.1) were commercially synthesized as full-length DNA copies (Biomatik GmbH, Rodgau, Germany) and supplied as plasmid clones. Viral complementary DNA (cDNA) sequences were PCR amplified from these vectors and subcloned into pDUAL plasmids using BsmB1 restriction endonuclease sites ^28^ such that the cDNA was flanked by bi-directional RNA polymerase I and RNA polymerase II promoters engineered to express viral (vRNA) and messenger RNA (mRNA), respectively. To generate segment 7 variants, mutations at nucleotide positions 52 (G->C) and 145 (G->A) to destroy the splice donor sequences for mRNAs 2 and 4 respectively were introduced into the segment cDNAs by oligonucleotide-directed mutagenesis (Figure 1A). Recombinant viruses Ck/Penn/G52C (ΔM2^-^/M42^+^) or Ck/Penn/G145A (M2^+^/ΔM42^-^) were rescued on *in-vitro* co-cultures of DF-1 and HEK 293T cells (1:1) as previously described ^28^; either as 7:1 reassortants with segment 7 of the Ck/Penn virus on a PR8 background (for BSL2 work) or as fully avian viruses handled at BSL3.

For vaccine challenge studies, the second egg passage was used for homologous challenge with A/chicken/Pennsylvania/1370/1983 H5N2 (Ck/Penn/83), or heterologous challenge with A/chicken/Queretaro/14588-19/1995 H5N2 (Ck/Queretaro/95), A/Hong Kong/483/1997 H5N1 (A/Hong Kong/97) and A/Northern Pintail/Washington/40964/2014 H5N2 (NoPT/Washington/14). Growth of the rg viruses was compared on DF-1 cells as previously described ^29^. The fully avian viruses were propagated and back titrated in specific-pathogen-free (SPF) embryonated chicken eggs (ECE) according to standard procedures ^30^. PR8-based viruses were propagated in ECE and titrated in MDCK cells. For partial purification of released virions, allantoic fluid was cleared of cell debris by centrifugation at 6,500 *g* for 10 minutes and 2.2 ml of the allantoic fluid was then overlaid over 1.1 ml of 30% sucrose in PBS and centrifuged at 125,000 *g* for 90 minutes. Virus pellets were then resuspended in Laemmli sample buffer for SDS-PAGE analysis.

### Protein and microscopy analyses

For western blotting, protein samples were denatured in Laemmli sample buffer, separated by SDS-PAGE and transferred to nitrocellulose membranes according to standard protocols. Rabbit polyclonal antisera to PR8 NP (2915) has been previously described ^31^, as has rabbit antiserum raised against whole PR8 virus ^32^. A goat antiserum raised against the whole M2 protein (G74) was the kind gift of Professor Alan Hay of the World Influenza Centre, London. An anti M42 ectodomain (M42e) serum was raised in rabbits to a synthetic peptide NH_2_-MSLQGKTLILRLTRNGWEC conjugated to Keyhole limpet haemocyanin (Genosphere Biotechnologies). A mouse monoclonal antibody (14C2) reactive against the M2 ectodomain (M2e) of PR8 was purchased from Abcam. Bound antibodies were detected using secondary anti-IgG antibodies conjugated to far-red fluorophores (Li-Cor) using a LiCor Odyssey imager.

For confocal microscopy, cells seeded on glass coverslips were washed in PBS and fixed with 4% formaldehyde in PBS for a maximum of 20 minutes, then washed three times with PBS/1% FBS before staining with anti-PR8 NP, followed by anti-rabbit IgG conjugated to Alexa-fluor 488 (Thermo Fisher). Coverslips were then mounted on glass slides using ProLong Gold anti-fade mountant containing DAPI and imaged using a Zeiss LSM710 confocal microscope and a 63x oil objective lens. Images presented in this study are maximum intensity projections of Z-stacks taken across the depth of the plasma membrane at approximately 0.30-0.45 μm increments. To quantify the filamentous budding phenotype two measurements were collected: the percentage (%) of infected cells producing filamentous bundles, and the average length of these bundles. Virus-infected cells were imaged and a total of 6 pictures portraying different fields of view were collected per virus over a total of three independent experiments. The % of infected cells producing filaments was then calculated. To determine filament length, 15 filamentous bundles were randomly selected per field of view and their length measured in ImageJ ^33^ in pixels and converted to μm as indicated: bundle length (pixels) × scale bar (pixels) / 20 μm (the scale bar length).

### Animals

For all experiments, mixed-sex SPF white leghorn chickens were obtained from and housed at the SEPRL at biosafety level 3 (BSL3E). Birds were maintained in HEPA filtered isolation cabinets with access to feed and water *ad libitum*. All experimental bird procedures were approved and performed under the SEPRL IACUC.

### Infection and transmission of wild-type, G52C, G145A viruses-Experiment 1

At 40 days of age, 3 groups of SPF chickens (n=5 per group) were infected via the intraocular/intranasal (IO/IN) route with 10^6.0^ EID_50_ of the following avian viruses: Ck/Penn/, Ck/Penn/G52C, or Ck/Penn/G145A in a volume of 0.2 ml (1/2 per route). On day 1 post-infection, two additional birds of the same age were placed in each isolator to assess the transmission of each virus to naïve-contact birds. All chickens were observed daily for development of clinical signs of disease and mortality. Oropharyngeal (OP) and cloacal (CL) swabs were collected from all birds from days 1 to 6 post-inoculation to assess infection/shedding loads, and transmission of the virus to naïve-contact birds. Swabs were placed into 2 ml of sterile brain-heart-infusion (BHI) supplemented with antibiotics as previously described ^19^. Birds were bled for sera collection at days 0 and 14 post-inoculation to examine seroconversion and euthanized at day 14 post inoculation.

### Vaccine protection with wild-type, G52C and G145A viruses from HPAI challenge-Experiment 2

At 28 days of age, groups of SPF chickens (n=10 per group) were vaccinated via IO/IN route with 10^6.0^ EID_50_ of either Ck/Penn, Ck/Penn/G52C, or Ck/Penn/G145A virus administered in a volume of 0.2 ml (1/2 per route). Sham vaccinated birds (n=10 per group) received PBS via the same volume and route. At 42 days of age (2 weeks post-vaccination), all chickens were bled and challenged with 10^6^ EID_50_ of either Ck/Penn/83 H5N2, Ck/Queretaro/95 H5N2, Ck/Hong Kong/97 H5N1 or NoPT/Washington/14 H5N2. Birds were monitored daily for mortality and clinical signs of disease. All birds were swabbed via oral and cloacal routes on days 2 and 4 post-challenge for detection of virus shedding. Swabs were placed into 2 ml of sterile brain-heart-infusion (BHI) as above. At 56 days of age (2 weeks post-challenge) all remaining birds were bled and euthanized.

### Comparison of vaccine protection using inactivated wild-type, G52C and G145A as vaccine virus with HPAI challenge-Experiment 3

Monovalent WT, G52C and G145A vaccine viruses were prepared following growth in SPF eggs and inactivated with beta-propiolactone (BPL) as previously described ^34^. At 21 days of age, four groups of SPF white leghorns (n=12 birds per groups) were vaccinated with one of the inactivated viruses: Ck/Penn, Ck/Penn/G52C, Ck/Penn/G145A or a sham-vaccine. Inactivated vaccines were composed of Montanide (70/30) at 512 hemagglutinating units (HAU) per dose and were administered subcutaneously in the nape of the neck in a total volume of 0.2 ml per bird. Sham birds received adjuvant containing virus-negative allantoic fluid. At 42 days of age (3 weeks post vaccination), serum was collected from all the birds to assess humoral immunity following vaccination and birds were challenged with 10^6^ EID_50_ of Ck/Penn/83 HPAIV. At 56 days of age, all surviving chickens were bled for collection of sera and were euthanized.

### Detection of virus from swab samples

Viral RNA extraction was carried out from OP and CL swabs with the MagMAX-96 AI-ND viral RNA isolation kit (Ambion, Austin, TX, USA) using an automated KingFisher™ magnetic particle processor (Thermo Fisher Scientific, Pittsburgh, PA, USA) as previously described ^35^. Primers and probe targeting the AIV matrix gene ^36^ were used to perform the quantitative RT-PCR (qRT-PCR) reaction by using the One-Step RT-PCR Kit (QIAGEN, Valencia, CA, USA) on a StepOne Real-time PCR System (Applied Biosystems, Darmstadt, Germany). For virus quantification, a standard curve was established with RNA extracted from dilutions of the same titrated stock of the viruses to convert data to EID_50_ equivalents. Cycle threshold (Ct) values of each viral dilution were plotted against viral titers. The resulting standard curve had a high correlation coefficient (R^2^ > 0.99; data not shown) and was used to convert Ct values to EID_50_/ml equivalents.

### Serology

Serum was obtained from all birds on days 0 and 14 post-vaccination or challenge and tested by hemagglutination inhibition (HI) assay using chicken red blood cells (USDA, SEPRL). The HI assay was performed using inactivated Ck/Penn/83 or Ck/Queretaro/95 HPAIV as antigens according to standard procedures ^37^. Titers were calculated as the highest reciprocal serum dilution providing complete hemagglutination inhibition. Serum titers of 1:8 (3 log_2_) or higher were considered positive for antibodies against AIV.

### Phylogenetic analysis

AIV HA sequences were obtained from GenBank and the Influenza Research Database (http://www.fludb.org/brc/home.do?decorator=influenza). Sequences were aligned with Clustal V (Lasergene 10.0, DNAStar, Madison, WI), and phylogenetic analysis for the H5 HA gene was performed with MEGA version 7 ^38^.

### Electron Microscopy

Viruses were BPL-inactivated (as above) and adsorbed onto freshly glow-discharged 400 mesh carbon parlodion-coated copper grids (Poly-Sciences, Warrington, PA). The grids were rinsed with buffer containing 20 mM Tris, pH7.4, and 120 mM KCl and negatively stained with 1% phospho-tungstic acid, then dried by aspiration. Viruses were visualized and measured on a JEM-1210 transmission electron microscope (JEOL USA, Inc., Peabody, MA) at an operating voltage of 120 kV at the University of Georgia. A minimum of 50 virions per group were measured for size comparisons of length and width.

### Statistical analysis

Comparisons of HI antibody titers between groups were performed by using one-way ANOVA with Tukey’s multiple comparison test for comparisons of all group means. Viral shedding infectious titers were compared between groups over time using one-way ANOVA with Tukey’s multiple comparison test as above, P ≤ 0.05 was considered statistically significant. All graphs and statistical tests were conducted using GraphPad Prism software version 7.0 (GraphPad Software Inc., San Diego, CA, USA).

## Results

### Characterization of wild-type, G52C and G145A viruses

Previously, we demonstrated that segment 7 from an HPAIV isolate from the 1983 Pennsylvania outbreak (A/Chicken/Pennsylvania/10210/1983) produced large amounts of mRNA4 and was therefore predicted to express a mixture of M2 and M42 ^6^. To probe the biological significance of this, we introduced mutations previously shown ^6^ to disrupt either the mRNA2 (G52C) or mRNA4 SD sequences (G145A) into segment 7 from the LPAIV progenitor of the Pennsylvania outbreak, A/chicken/Pennsylvania/1/1983 (Ck/Penn/83), to produce constructs expected to only express M42 (G52C) or M2 (G145A) respectively (Figure 1A). The 1983 Pennsylvania HPAIV outbreak was unusual in that the HA of the LPAIV progenitor possessed a polybasic cleavage site whose HPAIV phenotype was masked by glycosylation ^39, 40^. Accordingly, for initial characterization, wild-type and mutant segment 7s were rescued as 7:1 reassortants in the PR8 background, to permit work at BSL2. All viruses rescued readily and grew to high titers in ECE with no significant difference in titer levels (Figure 2A). To examine the effects of the splice site mutations on M2/M42 expression, virus particles were partially purified by pelleting through a sucrose pad before western blot analysis, using allantoic fluid from mock-infected eggs or eggs infected with WT PR8 as controls. No viral proteins were detected in the uninfected sample, while all infected samples contained similar amounts of the viral nucleoprotein (Figure 2B). The virus preparations also contained consistent amounts of viral ion channel, as assessed by using a goat polyclonal antiserum raised to full length M2, although it was noticeable that the PR8 polypeptide migrated faster than the Penn1 species. However, only the PR8 protein reacted with the M2 ectodomain (M2e)-specific monoclonal antibody 14C2, as expected given the sequence differences between PR8 and Penn1 ectodomains (Figure 1B) and the epitope specificity of the antibody ^41^. Conversely, only material from viruses with a Ck/Penn segment 7 reacted with an antiserum raised against the predicted Penn1 M42e sequence (Figure 2B). This demonstrated clear construct-dependent differences in relative staining intensity; lowest in the G145A virus and highest in the G52C virus, indicating that the mutations had successfully altered the antigenicity of the M2 ectodomain. This is consistent with the Ck/Penn segment producing both M2 and M42 and the splice site alterations producing M2 deficient (G52C) and M42 deficient (G145A) variants, as previously shown for A/Chicken/Pennsylvania/10210/1983 ^6^. Previous attempts to make polyclonal anti-peptide sera specific to the M42e sequence have shown residual cross-reactivity with M2e ^6^, presumably because of the common sequences at N- and C-terminal ends of the domain (Figure 1B). When virus growth was examined in two chicken cell lines (Figure 2A and B): the fibroblast line DF-1 ^42^ and the macrophage-like HD-11 line ^26^, Ck/Penn or Ck/Penn/G145A viruses grew similarly, but the Ck/Penn/G52C virus exhibited noticeably slower replication kinetics, without reaching statistical significance except for the comparison of Ck/Penn/G145 versus Ck/Penn/G52C in HD-11 cells (*p* = 0.002) (Figures 2C, D). Thus overall, altering the splice sites in Penn1 segment 7 modulated M2/M42 expression as expected and modestly altered virus growth in two chicken cell lines, but not embryonated eggs.

**Fig 2.**
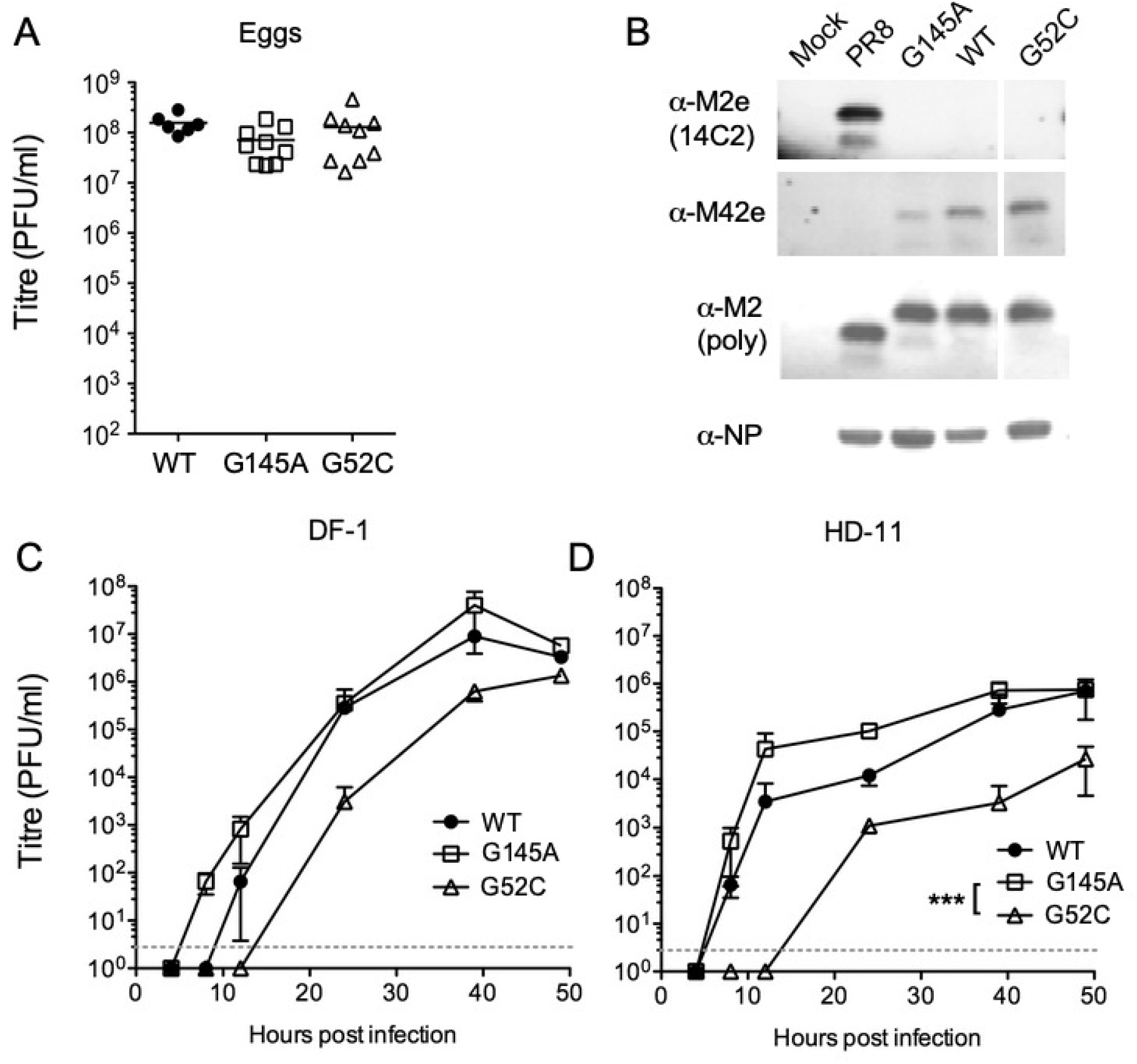
Characterization of rg PR8 7:1 reassortant viruses with Ck/Penn/83 WT, G52C and G145A segment 7s. (A) 11-day old embryonated eggs were inoculated with 1000 PFU of the indicated viruses before incubation at 37.5°C for two days. Allantoic fluid titers from individual eggs were determined by plaque assay. Bars indicate the mean. (B) Virus particles from eggs inoculated with the indicated viruses as in (A) or PBS (Mock) were partially purified by pelleting through a 30% sucrose pad before analysis by SDS-PAGE and western blotting with the indicated antisera. (C, D) DF-1 or HD-11 cells were infected at an MOI of 0.001 and virus titers measured at the indicated time points by plaque assay. The mean±S.D. of three independent experiments are plotted. The dashed line indicates the limit of detection. Growth curve data were analyzed by 2-way ANOVA and main effects model with Tukey’s post test comparison of virus groups. *** = p < 0.001.

Several reports have linked M2 expression and/or function to virus budding, both with regard to membrane scission and virus particle shape ^43–45^. To determine if altering the balance of M2 and M42 expression affected virus budding in this system, we examined the cell surface distribution of HA on infected cells by confocal microscopy using WT PR8 and a 7:1 reassortant with segment 7 from the human H3N2 A/Udorn/307/72 strain (PR8 MUd) as examples of viruses with non-filamentous and filamentous budding phenotypes respectively ^31, 46^. Mock infected cells gave only background staining, while cells infected with WT PR8 gave the expected stippled pattern of HA staining, indicative of non-filamentous virus budding (Figure 3A). In contrast, cells infected with PR8 MUd showed profuse bundles of viral filaments, many of which were over 20 µm long. Cells infected with the Ck/Penn or Ck/Penn/G52C reassortant viruses gave similar cell surface HA staining to WT PR8, but the Ck/Penn/G145A virus produced obvious bundles of short filaments. Quantification of replicate experiments indicated that the Ck/Penn/G145A filaments were on average 6 µm long and produced by around a third of the infected cells (Figures 3B, C). Thus, preventing M42 expression from Ck/Penn segment 7 introduced a moderately filamentous budding phenotype.

**Fig 3.**
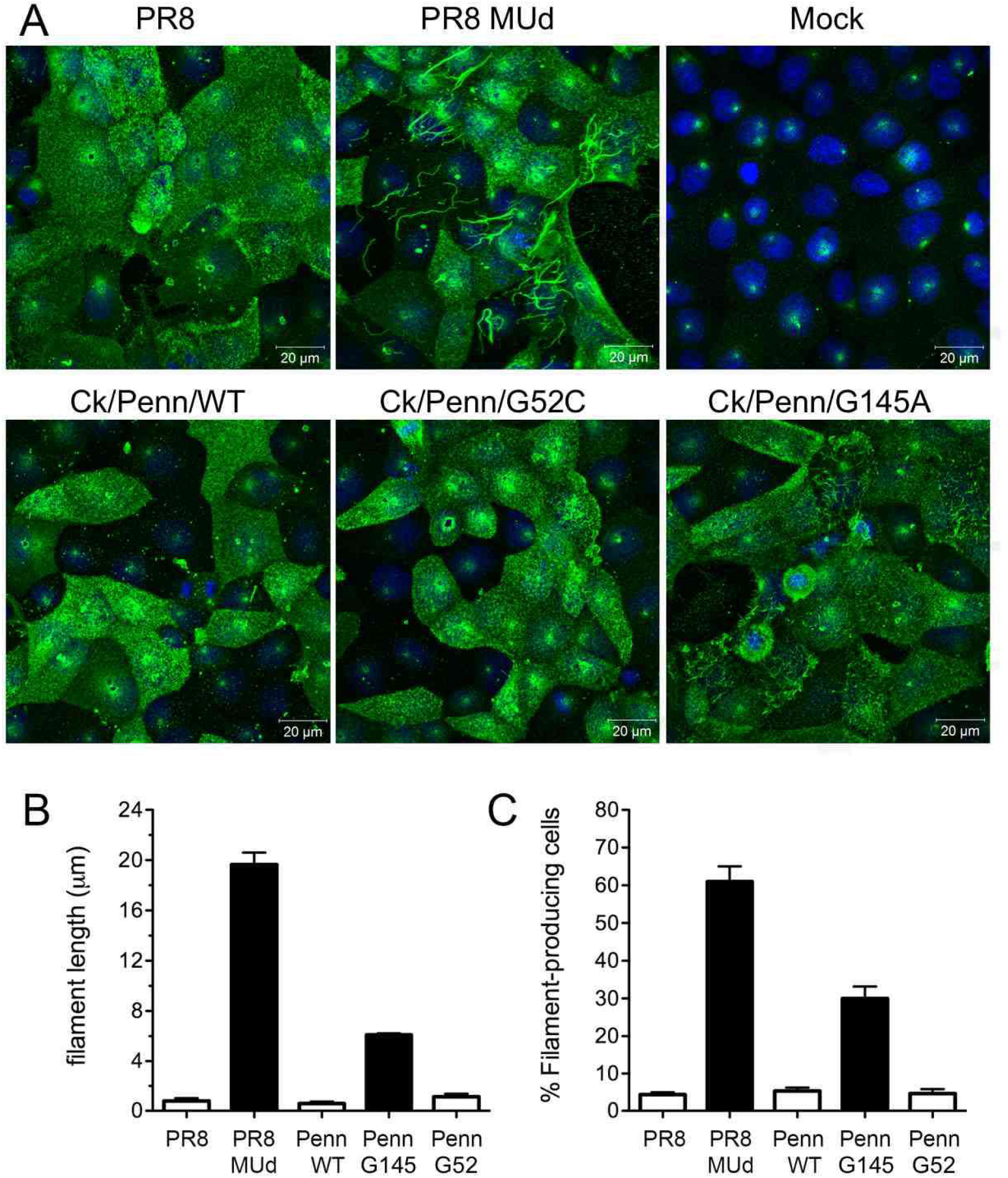
Budding phenotype of rg 7:1 reassortant viruses with Ck/Penn/83 WT, G52C and G145A segment 7s. MDCK cells were infected at an MOI of 3 with the indicated viruses, fixed at 16 h p.i. and surface-stained with anti-PR8 serum (green) before counterstaining for nuclei with DAPI (blue) and imaging by confocal microscopy. Images are maximum intensity projections of z-stacks taken through the depth of the cell. (B) The lengths of up to 90 filament bundles (_μ_M) were measured using ImageJ. (C) The percentage (%) of filament producing cells. A minimum of 100 infected cells were scored according to whether viral filaments were present or not. Numerical data are mean ± S.D. of measurements taken from three independent experiments.

To further examine the consequences of altering M2 isoform expression on AIV biology, we attempted to re-rescue the mutant segment 7s in the background of the other seven Ck/Penn segments. For reasons that remain unclear, this was unsuccessful (data not shown). However, replacing Ck/Penn segment 1 with that of an H7N3 AIV (A/Anhui/1/2013) produced viable viruses. Following rescue of these rg viruses, the effects of altering M2 or M42 expression were assessed on virus growth on DF-1 cells, in comparison to the original non-recombinant Ck/Penn/WT virus. The growth kinetics of the rg viruses compared to the original virus were similar, reaching a plateau at 24 h.p.i., but the peak titers were variable, with the rg WT (hereafter, just WT) virus growing best and the G52C virus worst, plateauing around 1log_10_ lower (Figure 4A). To examine the effects of altering ion channel expression on virion morphology in this system, virus particles released into the allantoic cavity of infected eggs were examined by negative stain electron microscopy. The Ck/Penn/WT virus produced the expected roughly spherical virions with average major and minor axis diameters of around 120 and 110 nm respectively, but viruses expressing only the M2 or M42 proteins showed much greater size variation, with the G145A virus showing obvious filamentous particles (Figure 4B, C). When considered as a ratio of major to minor axis length, Ck/Penn/G145A virus particles had a significantly different size characteristic to the other two viruses (Figure 4D), consistent with filamentous budding. Thus, altering M2/M42 expression in an AIV genetic background also altered virus budding morphology.

**Fig 4.**
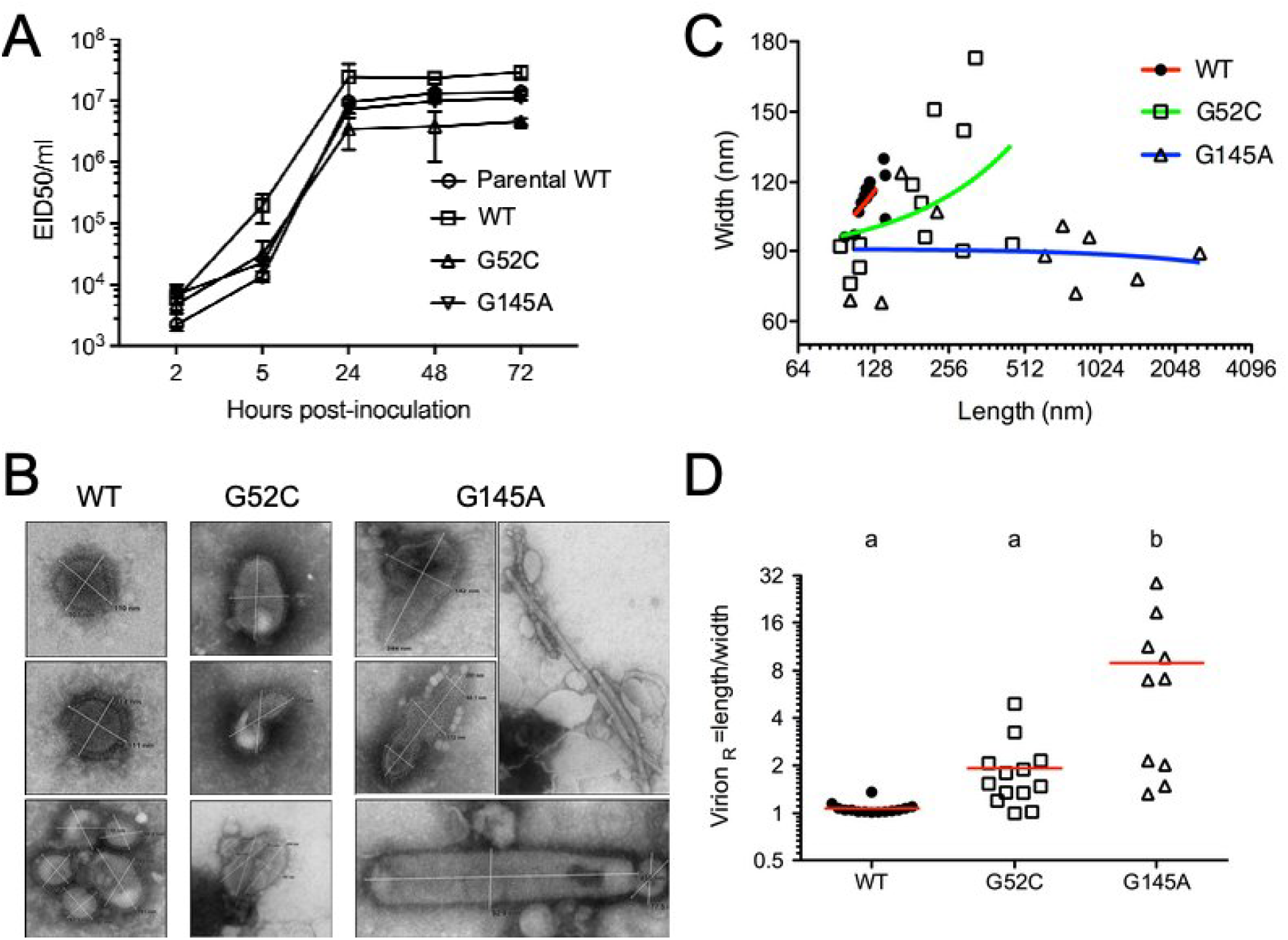
Characterization of Ck/Penn/83 viruses with WT, G52C and G145A segment 7s. (A) Growth kinetics in DF1 cells in EID_50_ titers. Cells were infected at an MOI of 0.1, harvested, and tittered at the indicated time points. Data are the mean ± SD of three independent experiments. (B) Viruses were grown in 9-11 day old SPF chicken eggs, collected from allantoic fluid, and subjected to negative stain electron microscopy with 1 % phosphor-tungstic acid (C). Virion size was characterized by measuring the major and minor axis length of a minimum of 50 particles. Linear regression analysis was performed in GraphPad Prism. (D) Virion ellipticity was determined by taking the ratio of long and short axes. Lines indicate the mean virion length versus width ratio (R). Groups with different subscript letters are significantly different (p<0.05).

### The Ck/Penn/G52C and Ck/Penn/G145A viruses replicate *in vivo* but do not transmit

In mammalian models of influenza A virus infection, virion morphology has been suggested to influence transmission ^47–49^, but this has not been tested for AIV. To determine infection and transmissibility of the Ck/Penn/WT, Ck/Penn/G52C, and Ck/Penn/G145A viruses, groups of SPF chickens were inoculated via IO/IN with each virus and two further birds were introduced into each of the isolators one day later. None of the inoculated or contact birds demonstrated clinical signs of diseases or mortality during the experiment (data not shown). However, all three viruses were able to infect chickens, as viral RNA was detected from days 1 to 6 post-inoculation in oral swabs (Figure 5A) but not in cloacal swabs (data not shown). Titers (estimated from the qRT-PCR values using a standard curve derived from the same virus stocks) of the viruses were similar among infection groups on days 1 to 6 post-inoculation. Introduction of susceptible birds into each of the experimental groups resulted in the transmission of the wild-type virus, with similar levels of virus replication in both contact birds (Figure 5A). However, transmission of the Ck/Penn/G52C and Ck/Penn/G145A viruses was not observed, as all swabs were negative from contact birds.

**Fig 5.**
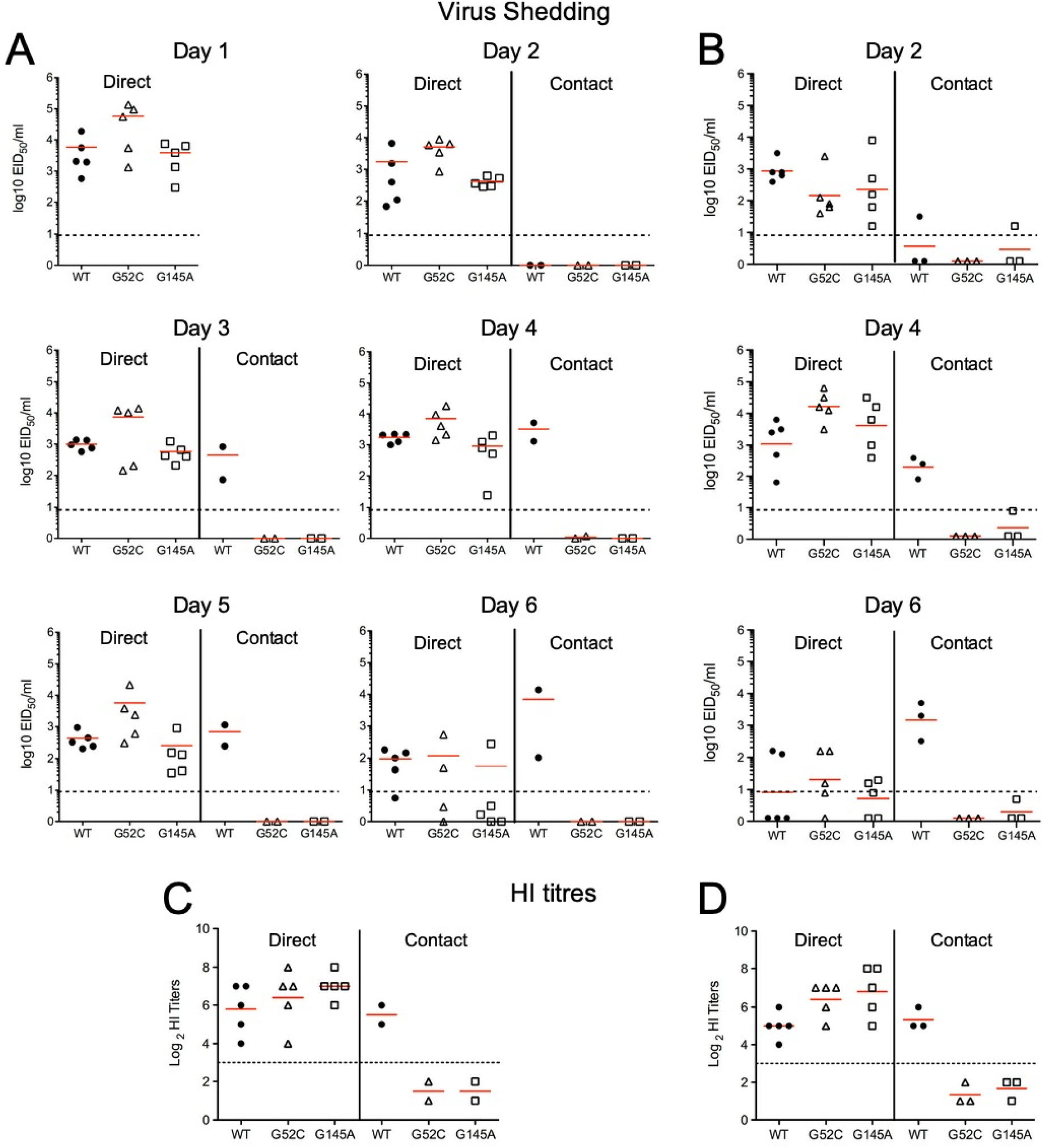
Shedding, transmission and humoral immune responses of the Ck/Penn, Ck/Penn/G52C and Ck/Penn/G145A segment 7 mutant viruses. (A) Directly inoculated chickens (n=5) received 10^6^ EID_50_ per bird of the viruses on day 0. On day 1 post-inoculation, two contact birds were placed in each unit. Oral swabs were collected daily for 6 days and virus load determined by qRT-PCR. (B) Birds were bled on day 0 and 14 post-inoculation and antibody titers to Ck/Penn/83 determined by HI assay. (C) Directly inoculated chickens (n=5) received 10^6^ EID_50_ per bird of the viruses on day 0. On day 1 post-inoculation, three contact birds were placed in each unit. Oral swabs were collected on days 2, 4, and 6 post-inoculation and virus load determined by RT-PCR. (D) Birds were bled on day 0 and 14 post-inoculation and antibody titers to Ck/Penn/83 determined by HI assay. Dashed lines indicate the minimum titer considered as positive (3log_2_). Red lines indicate the mean.

To determine if the failure of the G52C and G145A viruses to transmit was reproducible, the experiment was repeated using a larger cohort of contact birds added at day 1 p.i. As before, all directly inoculated birds secreted readily detectable amounts of virus from the oral cavity at days 2 and 4 p.i. but were beginning to clear the virus by day 6, with no sign of attenuated replication of the mutant viruses (Figure 5B). Once again, the Ck/Penn/WT virus transmitted efficiently to the contact birds, and while one contact bird sporadically shed very low levels of the Ck/Penn/G145A virus, no detectable transmission was observed in birds infected with the Ck/Penn/G52C virus. Finally, HI testing was performed on all birds to determine if seroconversion occurred as a confirmatory indicator of infection. The results demonstrated that in both transmission studies all birds directly infected with virus seroconverted after 2 weeks (Figures 5C, D). In contrast, only contact birds in Ck/Penn/WT groups seroconverted to AIV, confirming transmission of that virus to naïve cohorts. However, neither of the segment 7 variants, G52C or G145A, were able to transmit to contact birds based on a lack of seroconversion, supporting the virus shedding data. Thus, the Ck/Penn/G52C and Ck/Penn/G145A viruses replicated in directly inoculated birds but showed no, or very limited transmission to contact birds.

### The Ck/Penn/G52C and Ck/Penn/G145A live virus vaccines protect against homologous and heterologous HPAI challenge

Vaccine efficacy was assessed by measuring survival and viral shedding from chickens infected (or sham infected) with the LPAIV wild-type or mutant rg viruses and challenged 2 weeks later with either the homologous H5N2 HPAIV Ck/Penn/1370 or three increasingly antigenically– distant heterologous H5 HPAIV. The heterologous HPAIV used in these studies were: Ck/Queretaro/1995 (H5N2) from the same North American lineage as Ck/Penn/1370, sharing approximately 90 % sequence similarity of HA; the Asian lineage clade 0 H5N1 A/Hong Kong/97 virus (∼ 74% similarity) and an H5N2 clade 2.3.4.4b isolate NoPT/Washington/14 (∼70% similarity) (Figure 6). All sham-vaccinated birds died following challenge, with mean death times of 5.5, 2.0, 6.5 and 4 days against Ck/Penn/1370, Ck/Queretaro/95, A/Hong Kong/97 and NoPT/Washington/14, respectively (Figure 7). In contrast, all birds that received the rg LPAIV vaccine viruses, whether containing a WT or mutant segment 7 survived lethal challenge with both homologus and heterologous HPAIV. No clinical signs of disease were observed in any of the vaccinated and challenged birds. Thus, the live virus H5N2 vaccines provided complete protection against a wide range of antigencally distant H5 challenge viruses.

**Fig. 6.**
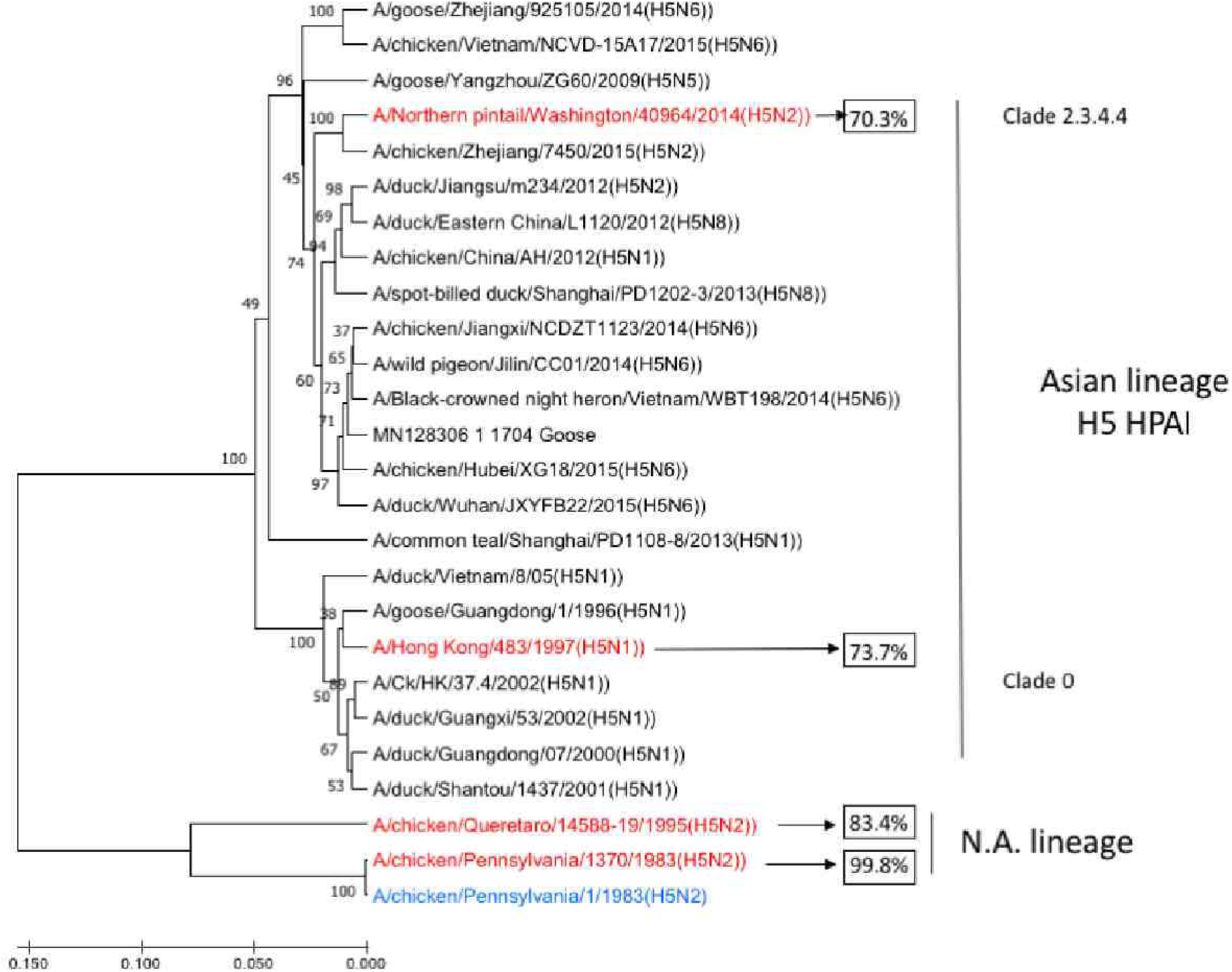
Evolutionary relatedness of selected H5 HAs. A phylogenetic tree of H5 HAs from the indicated viruses was constructed to show the degree of relatedness of the vaccine (blue) and challenge (red) viruses used in this study. Values in boxes give the % amino acid similarity between vaccine and challenge viruses.

**Fig 7.**
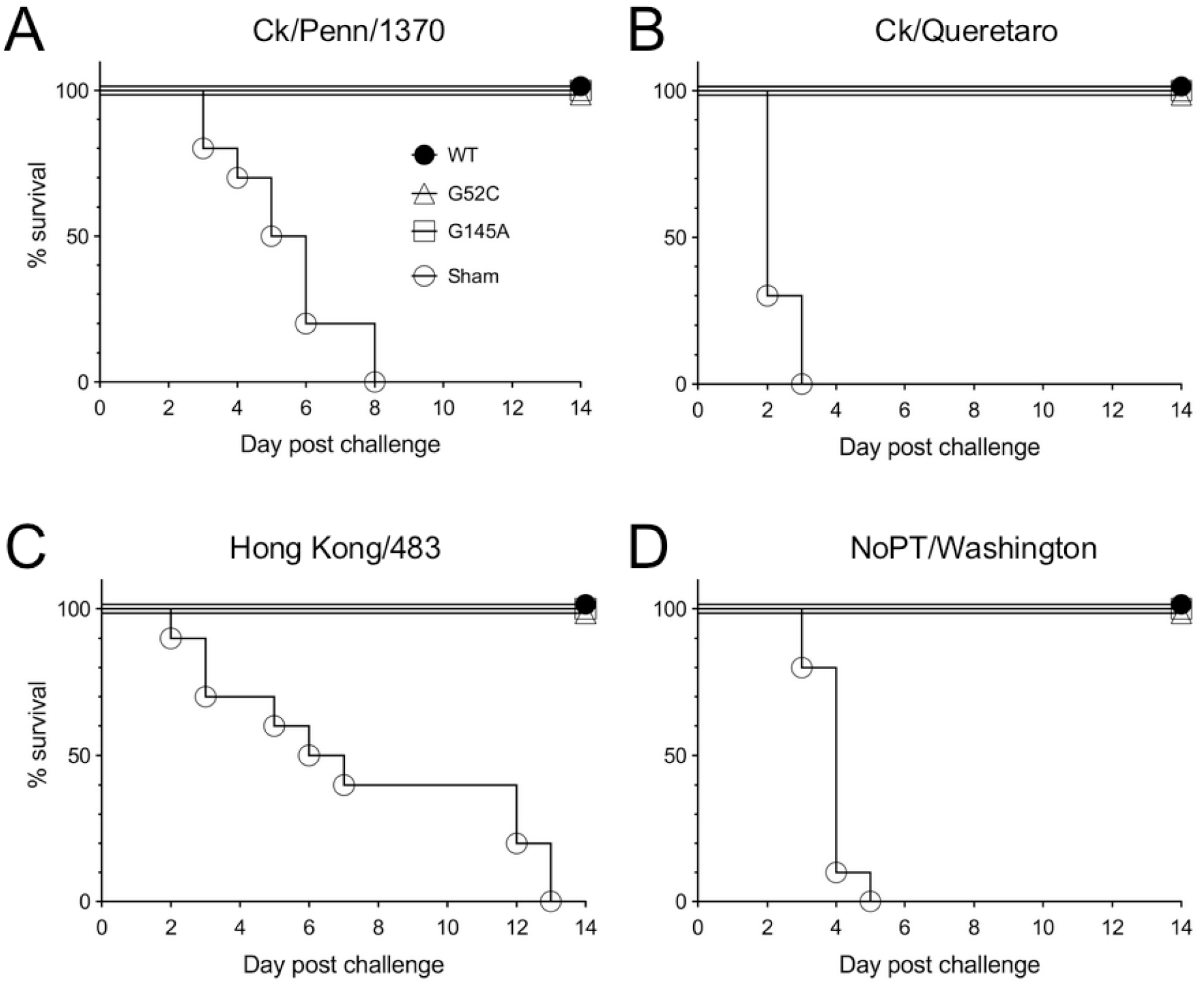
Vaccine efficacy trials with Ck/Penn and segment 7 mutant viruses. Groups of 4-week-old birds were vaccinated with 10^6.0^ EID_50_ of either Ck/Penn WT, Ck/Penn/G52C, Ck/Penn/G145A virus or sham vaccinated with PBS. Two weeks later, all birds were challenged with 10^6.0^ EID_50_ of the indicated HPAIV. Kaplan-Meier survival plots are shown.

High levels of HPAIV virus shedding were observed in the sham-vaccinated birds for all challenge viruses, from both OP and CL routes, with maximum values ranging from approximately10^6^ EID_50_/ml on day 4 post challenge with NoPT/Washington to 10^3^ EID_50_/ml on day 2 post challenge with Hong Kong/483 (Figure 8). In contrast, virus shedding was significantly reduced in all groups of birds vaccinated with either of the Ck/Penn, Ck/Penn/G52C and Ck/Penn/G145A live virus vaccines, although not completely blocked in all birds in all instances.

**Fig 8.**
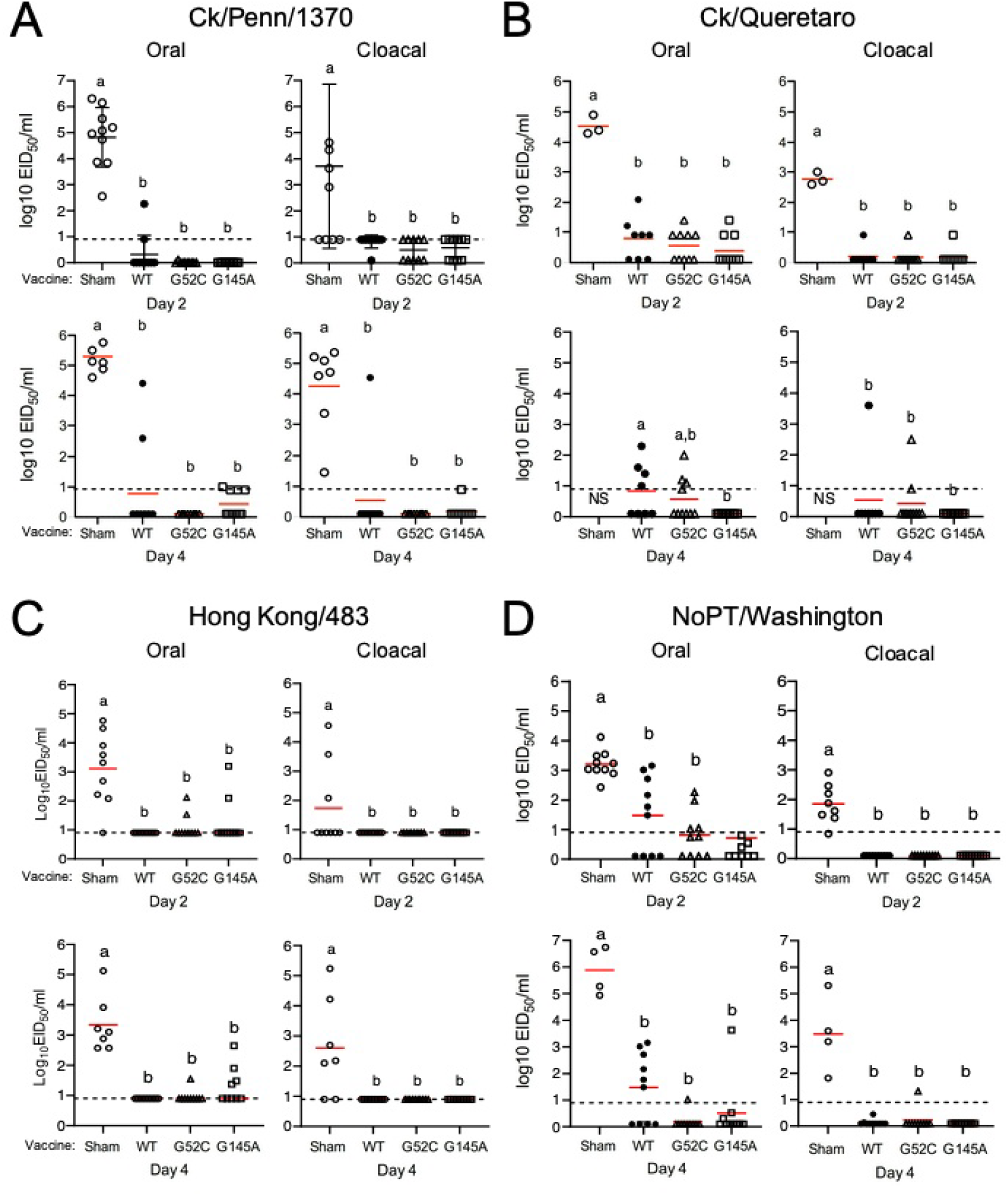
HPAIV shedding by vaccinated and unvaccinated birds. Birds were vaccinated (or sham vaccinated) with Ck/Penn, Ck/Penn/G52C or Ck/Penn/G145A live virus vaccines at 4 weeks-of-age, and challenged at 6 weeks-of-age with HPAIV. Challenge viruses used were Ck/Penn/83 (A), Ck/Queretaro/95 (B), A/Hong Kong/97 (C), and NoPT/Washington/15 (D). Viral titers were determined from oral and cloacal swabs taken on days 2 and 4 post-challenge. The lower limit of detection (LOD) was 0.9 log_10_ EID_50_ per ml.

To examine the level of cross reactive antibodies induced following vaccination and challenge, HI titers against the homologous (Ck/Penn/1370) and heterologous (Ck/Queretaro/95) challenge HA antigens were determined. In the case of homologous challenge, all groups of vaccinated birds exhibited high HI titers to the challenge HA two weeks post vaccination of approximately 6-7 log_2_ to the homologous antigen, which did not significantly increase following challenge when the homologous vaccine antigen was used in the assay (Figure 9A). When these same sera were tested against the heterologous antigen (Queretaro), the titers dropped by approximately 3log_2_ (Figure 9B). Serum collected following challenge with the heterologous Ck/Queretaro HPAIV was also tested against both antigens. After vaccination and challenge, HI titers increased significantly using both antigens from around 4 log_2_ to 8 log_2_; however, no significant differences within the different vaccine groups were observed (Figures 9C and D). Thus the Ck/Penn/G52C and Ck/Penn/G145A vaccine viruses induced protective responses indistinguishable from the wild-type parental virus, despite their attenuated shedding phenotype.

**Fig 9.**
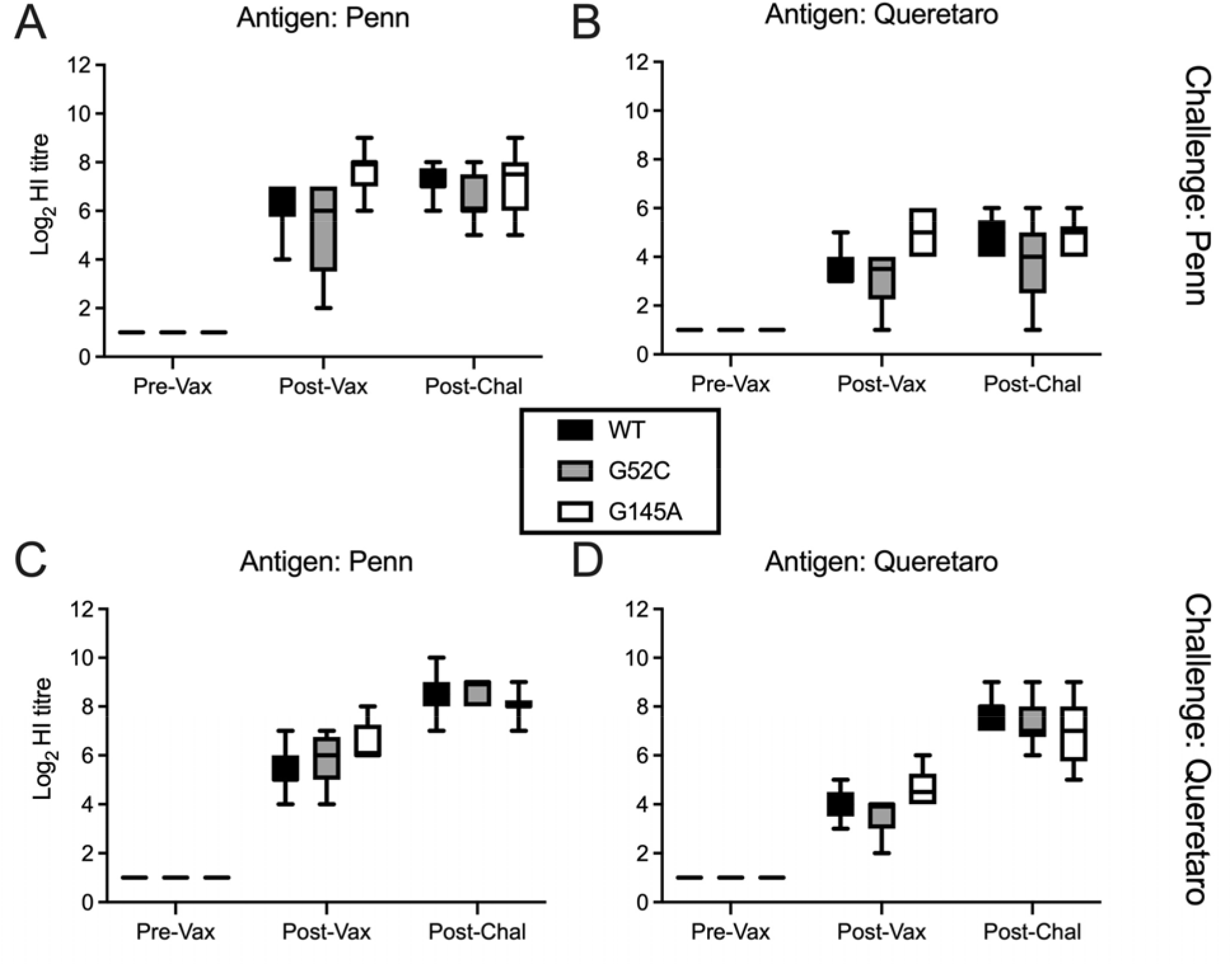
Cross reactive HI antibody responses from birds receiving live H5N2 vaccine and challenged with either homologous (Penn) or heterologous (Queretaro) HPAIV. Birds were vaccinated (or sham vaccinated) with Ck/Penn, Ck/Penn/G52C or Ck/Penn/G145A live virus vaccines at 4 weeks-of-age, and challenged at 6 weeks-of-age with the indicated viruses. Sera taken at 0 days pre-vaccination (Pre-Vax), 14 days post-vaccination (Post-Vac) and 14 days post-challenge (Post-Chal) respectively, were tested for reactivity against the indicated viruses. All sera from Sham vaccinated birds were ≤ 2log_2_ and are not shown. Titers are expressed as log_2_ geometric mean values. Samples with titers below 3log_2_ GMT were considered negative.

### Comparison of protective efficacy with inactivated Ck/Penn/G52C and Ck/Penn/G145A virus vaccines against homologous HPAI challenge

An accepted advantage of live-attenuated vaccines over inactivated ones is the induction of a broader response, including T-cells and mucosal immunity. To establish whether this held true for the Ck/Penn/G52C and Ck/Penn/G145A viruses, we tested the efficacy of vaccine protection using BPL-inactivated rg viruses in chickens following virus challenge with Ck/Penn/1370 HPAIV. Birds in this study received a single subcutaneous injection of 512 HAU of inactivated virus in oil emulsion adjuvant. All birds receiving inactivated wild-type, G52C, or G145A virus vaccines survived lethal challenge. (Fig. 10A). No clinical signs of disease were observed in any of the vaccinated-challenged birds. In contrast, all sham-vaccinated birds died following challenge, with a mean death time of 5.5 days. Serum was collected before and after vaccination and challenge for antibody testing by HI. Three weeks post vaccination HI titers were demonstrated to be approximately 9 log_2_ (Figure 10B), consistent with the three groups receiving the same HA antigenic load per dose. Following Ck/Penn/1370 challenge, the HI titers increased, but not significantly. As expected, oral swabbing showed that sham-vaccinated birds shed considerable quantities of virus (between 10^6^-10^7^ EID_50_/ml) via the oral route on days 2 and 4 post challenge (Fig 10C, E). Vaccinated birds shed significantly less (between 10^3^-10^4^ EID_50_/ml), but still readily detectable quantities of virus. This is in contrast to the performance of the live versions of the vaccines, where oral shedding from most animals was below the limit of detection (Fig 8A). Performance of the live (Fig 8) and killed versions of the vaccines in blocking cloacal shedding was similar, with the killed vaccines also taking shedding down to near undetectable levels (Fig 10 D, F). Thus while both live and inactivated vaccines provided complete disease protection from homologous HPAIV challenge, the live vaccines performed substantially better at decreasing virus shedding, which would result in decreased transmission potential to other birds. Overall the live virus vaccine reduced virus shedding 500-10,000-fold more than the inactivated vaccine (Fig 8A and Fig 10C-F).

**Fig 10.**
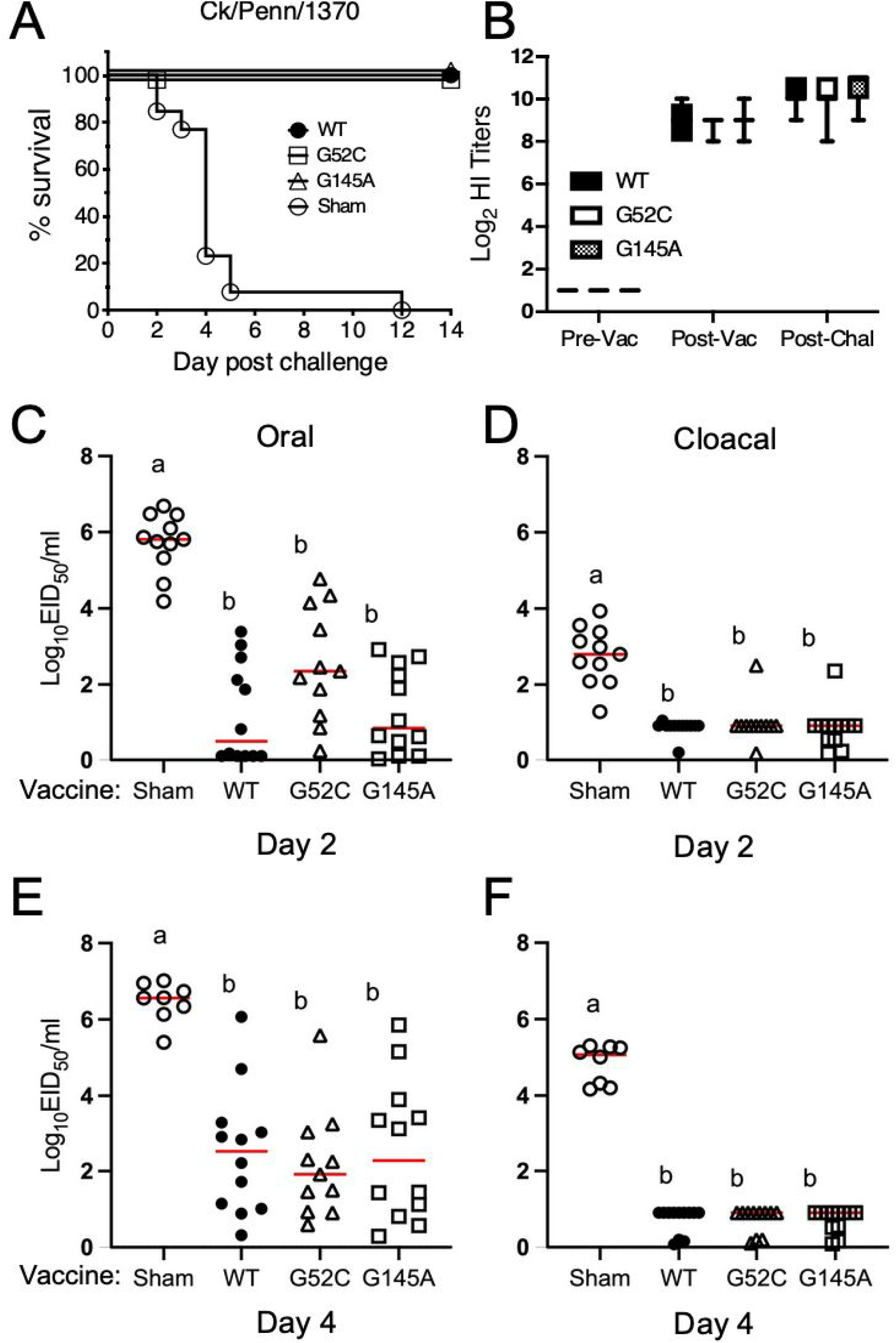
Protection of chickens using inactivated vaccines produced from the live H5N2 Ck/Penn viruses. Chickens were vaccinated with 512 HUA of adjuvanted BPL-inactivated Ck/Penn, Ck/Penn/G52C or Ck/Penn/G145A at 3 weeks-of-age and challenged 10^6^ EID_50_ per bird at 6 weeks-of-age with Ck/Penn/83/H5N2 HPAIV. (A) Kaplan-Meier survival plot following vaccination and challenge. (B) Serum HI titers from groups of vaccinated birds at pre / post vaccination and post challenge. All sera from Sham vaccinated birds were < 2log_2_ and are not shown. (C, D, E, F) Virus shedding titers following vaccination and challenge with Ck/Penn/83/H5N2 HPAIV. Titers are expressed as log_2_ GMT. Samples with titers below 3log_2_ GMT were considered negative. Viral titers from oral and cloacal swabs on day 2 (C, D) and 4 (E, F) post-challenge. Viral titers are expressed as log_10_ EID_50_ per ml. The lower limit of detection (LOD) was 0.9 log_10_ EID_50_ per ml.

## Discussion

Following the emergence of Goose/Guangdong lineage H5N1 in China in 1996, H5 HPAIVs have spread world-wide and continue to cause severe morbidity and mortality in domestic poultry, as well as sporadic zoonotic infections. Due to the genetic and antigenic diversity of these viruses, many Southeast Asian countries have incorporated AIV vaccination into their control programs to reduce mortality, environmental shedding, transmission to other flocks, and reduce potential human exposure ^14, 50^. Faced with recent annual winter epidemics of H5NX HPAIV, many European countries are now considering vaccination of poultry (https://www.wattagnet.com/articles/45210-europe-pushes-for-acceptance-of-hpai-vaccination). However, the constant evolution of AIV in nature requires constant updating of vaccines to best-match potential outbreak strains. Mismatched vaccines to outbreak strains may impart partial immunity and allow asymptomatic circulation of the virus, resulting in continued spread and evolution of the virus ^9, 51^. As a result, new strategies are needed to develop AIV vaccines that significantly reduce viral shedding and thus decrease transmission.

In these studies, live virus vaccines were created by mutating segment 7 of the Ck/Penn/83 H5 strain so that it only allowed the expression of either M2 (G145A) or M42 (G52C) instead of both M2/M42 isoforms of the viral ion channel protein. All rg viruses were able to infect chickens, replicate and induce protective immune responses. However, the G52C and G145A viruses did not demonstrate transmission to naive-contact birds. The reasons for this failure to transmit remain to be determined, but disrupting M2 or M42 expression affected virion morphology, in the case of the G145A virus to the point of inducing filamentous budding. In mammalian influenza A viruses, viral filaments have been associated with increased viral transmission ^47–49^, but in this case, the filamentous virus was unable to transmit efficiently. Alternatively, HPAIV HAs can be more sensitive to premature flipping into the fusion conformation at low pH and thus have a higher dependency on M2 expression lowering the pH of the *trans-*Golgi apparatus ^52–55^. Our alterations here to viral ion channel expression may have reduced protection of the Ck/Penn/83 HA against low pH during intracellular migration through exocytic pathway and thereby produced virions with fewer fusion-competent HA molecules that are less infectious and thus reduce transmission. HAs with reduced stability might lead to a lower level of fusion due to a lack of sufficient numbers of fusion-competent molecules to efficiently support cooperative conformational changes within an HA trimer or between the multiple HA trimers that are likely needed for fusion. Alternatively, it is conceivable that fusion may need to take place in a particular endosomal compartment to proceed to the next step of the infection. In this case, the pH of HA activation may need to be closely matched to the pH of the proper (infection-permissive) vesicular compartment. Finally, other factors such as virus origin, species, inoculation dose and age can all affect shedding patterns of AIV ^56^. Additional studies are underway to better define this effect.

Both G52C (ΔM2) and G145A (ΔM42) vaccines protected against homologous and heterologous H5N2 challenge that demonstrated their potential as live virus vaccine. The results from the vaccine experiments agreed with previous studies demonstrating the advantage of the use of live vaccines over inactivated vaccines ^23, 57^. In the live virus vaccination - challenge experiment here (Experiment 2), virus shedding after HPAIV challenge from vaccinated birds was below the level of detection, demonstrating the combined effects of developing multiple arms of immunity, including adaptive (humoral and cellular) and likely mucosal. In contrast, while the killed vaccine reduced viral shedding after HPAIV challenge compared to sham vaccinated birds, there was still significant viral shedding to the environment that might allow silent circulation and transmission to other flocks if challenge occurred in the field. The lack of protection against shedding might also be related to the lack of cellular and mucosal immunity induced by the parental application of the inactivated vaccine. In addition, killed vaccines do not follow the natural replication cycle of the virus and are not able to produce effective cell-mediated immunity, which is commonly associated with cross-protection against antigenically distant strains ^19, 58, 59^. Challenge with a heterologous virus resulted in complete protection against disease, and significantly reduced viral shedding. The broader mechanism of protection conferred by live vaccines is related to their mimicking of the natural infection and immunity against all the viral proteins and not just one viral target such as recombinant vaccines, and their eliciting of a broader immune response, including humoral and cell-mediated immunity. Administration of live viruses is more suitable and can be administered by spray or drinking water, reducing the cost of administration. It bears noting of the potential of LPAIV to shift, through either mutation or recombination, into a HPAIV. Concerns over this occurrence are a primary reason live AIV vaccines are not commercially used. One of the benefits demonstrated here is the lack of transmission through disruption of M2/M42. Whether this effect is maintained in other rg viruses is currently being studied as different HA subtype live vaccines are being tested.

The *in vitro* results replication studies demonstrated that IAV can utilize M2 or M42 segment as an ion channel interchangeably, which agrees with previous studies where mammalian PR8 virus was used ^6, 60^. Growth kinetics of the viruses in DF1 cells demonstrated that G52C (ΔM2) viruses had a disadvantage over the other viruses tested in terms of titer. Both the G52C and G145A (ΔM42) viruses demonstrated lower titers compared to the rg WT suggesting that both forms of ion channel are needed for optimal replication. Morphologically, influenza viral particles are pleomorphic, with typical spherical virions that are ∼100 nm in diameter and filamentous virions reaching 20 μm in length ^47–49^. Analysis of the morphology of the G52C and G145A viruses by electron microscopy demonstrated that viruses with altered M2/M42 expression were larger in size and had a filamentous form as compared to the viruses encoding just the M2 or M42 proteins.

In conclusion, we developed ΔM2 and ΔM42 vaccine viruses that can potentially be used as live vaccines in high-risk areas to not only reduce morbidity and mortality but also significantly reduce or abrogate virus replication following challenge with HPAIV in chickens. Currently the use of live AIV vaccines is not an alternative for the prevention of disease or infection. However it might be a suitable alternative in places where stamping out or the use of inactivated vaccines might not be suitable due to the lack of resources or the development of an efficient or matched vaccines because of the lack of a defined market.

## Acknowledgements

This study was supported by a US-UK BBSRC-NIFA Collaboration grant (no. 2015-67015-22968 (NIFA) and BB/M027163/1 (BBSRC)) to DRK, LV and PD. Additional funding was provided by USDA-ARS CRIS # 6040-32000-062-00D to DRK and BBSRC strategic program grant funding (BBS/E/D/20002173 and BBS/E/D/20002174) to LV and PD. Work described in this manuscript provides the basis for United States Patent no. WO/2020/185898 and European Patent no. EP3938496 “HA-specific influenza virus attenuated vaccine comprising mutations in segment 7, and uses therefor”, held by DRK, PD, LV and DLS.

## References

1 Dubois, J., Terrier, O. & Rosa-Calatrava, M. Influenza viruses and mRNA splicing: doing more with less. MBio 5, e00070–00014, doi:10.1128/mBio.00070-14 (2014).

2 Pinto, R. M., Lycett, S., Gaunt, E. & Digard, P. Accessory Gene Products of Influenza A Virus. Cold Spring Harb Perspect Med, doi:10.1101/cshperspect.a038380 (2020).

3 Lamb, R. A. & Choppin, P. W. Identification of a second protein (M2) encoded by RNA segment 7 of influenza virus. Virology 112, 729–737, doi:10.1016/0042-6822(81)90317-2 (1981).

4 Pinto, L. H. & Lamb, R. A. The M2 proton channels of influenza A and B viruses. J Biol Chem 281, 8997–9000, doi:10.1074/jbc.R500020200 (2006).

5 Shih, S. R., Suen, P. C., Chen, Y. S. & Chang, S. C. A novel spliced transcript of influenza A/WSN/33 virus. Virus Genes 17, 179–183 (1998).

6 Wise, H. M. et al. Identification of a novel splice variant form of the influenza A virus M2 ion channel with an antigenically distinct ectodomain. PLoS Pathog 8, e1002998, doi:10.1371/journal.ppat.1002998 (2012).

7 Suarez, D. L. & Senne, D. A. Sequence analysis of related low-pathogenic and highly pathogenic H5N2 avian influenza isolates from United States live bird markets and poultry farms from 1983 to 1989. Avian Dis 44, 356–364 (2000).

8 Capua, I. & Marangon, S. Control of avian influenza in poultry. Emerg Infect Dis 12, 1319–1324, doi:10.3201/eid1209.060430 (2006).

9 Kapczynski, D. R. et al. Homologous and heterologous antigenic matched vaccines containing different H5 hemagglutinins provide variable protection of chickens from the 2014 U.S. H5N8 and H5N2 clade 2.3.4.4 highly pathogenic avian influenza viruses. Vaccine 35, 6345-6353, doi:10.1016/j.vaccine.2017.04.042 (2017).

10 European Food Safety, A. et al. Avian influenza overview December 2021-March 2022. EFSA J 20, e07289, doi:10.2903/j.efsa.2022.7289 (2022).

11 Bevins, S. N. et al. Intercontinental Movement of Highly Pathogenic Avian Influenza A(H5N1) Clade 2.3.4.4 Virus to the United States, 2021. Emerg Infect Dis 28, 1006-1011, doi:10.3201/eid2805.220318 (2022).

12 Alexander, D. J. An overview of the epidemiology of avian influenza. Vaccine 25, 5637–5644, doi:10.1016/j.vaccine.2006.10.051 (2007).

13 Lycett, S. J., Duchatel, F. & Digard, P. A brief history of bird flu. Philos Trans R Soc Lond B Biol Sci 374, 20180257, doi:10.1098/rstb.2018.0257 (2019).

14 Richard-Mazet, A., Goutebroze, S., Le Gros, F. X., Swayne, D. E. & Bublot, M. Immunogenicity and efficacy of fowlpox-vectored and inactivated avian influenza vaccines alone or in a prime-boost schedule in chickens with maternal antibodies. Vet Res 45, 107, doi:10.1186/s13567-014-0107-6 (2014).

15 Kapczynski, D. R. & Swayne, D. E. Influenza vaccines for avian species. Curr Top Microbiol Immunol 333, 133–152, doi:10.1007/978-3-540-92165-3_6 (2009).

16 Wei, X. & Cui, J. Why were so few people infected with H7N9 influenza A viruses in China from late 2017 to 2018? Sci China Life Sci 61, 1442–1444, doi:10.1007/s11427-018-9406-4 (2018).

17 Suarez, D. L. & Pantin-Jackwood, M. J. Recombinant viral-vectored vaccines for the control of avian influenza in poultry. Vet Microbiol 206, 144–151, doi:10.1016/j.vetmic.2016.11.025 (2017).

18 Hghihghi, H. R. et al. Characterization of host responses against a recombinant fowlpox virus-vectored vaccine expressing the hemagglutinin antigen of an avian influenza virus. Clin Vaccine Immunol 17, 454–463, doi:10.1128/CVI.00487-09 (2010).

19 Kapczynski, D. R. et al. Vaccine protection of chickens against antigenically diverse H5 highly pathogenic avian influenza isolates with a live HVT vector vaccine expressing the influenza hemagglutinin gene derived from a clade 2.2 avian influenza virus. Vaccine 33, 1197–1205, doi:10.1016/j.vaccine.2014.12.028 (2015).

20 Niqueux, E., Guionie, O., Amelot, M. & Jestin, V. Prime-boost vaccination with recombinant H5-fowlpox and Newcastle disease virus vectors affords lasting protection in SPF Muscovy ducks against highly pathogenic H5N1 influenza virus. Vaccine 31, 4121–4128, doi:10.1016/j.vaccine.2013.06.074 (2013).

21 Steensels, M. et al. Prime-boost vaccination with a fowlpox vector and an inactivated avian influenza vaccine is highly immunogenic in Pekin ducks challenged with Asian H5N1 HPAI. Vaccine 27, 646–654, doi:10.1016/j.vaccine.2008.11.044 (2009).

22 Swayne, D. E. et al. Antibody titer has positive predictive value for vaccine protection against challenge with natural antigenic-drift variants of H5N1 high-pathogenicity avian influenza viruses from Indonesia. J Virol 89, 3746–3762, doi:10.1128/JVI.00025-15 (2015).

23 Kapczynski, D. R. et al. Protection of commercial turkeys following inactivated or recombinant H5 vaccine application against the 2015U.S. H5N2 clade 2.3.4.4 highly pathogenic avian influenza virus. Vet Immunol Immunopathol 191, 74-79, doi:10.1016/j.vetimm.2017.08.001 (2017).

24 Capua, I. & Marangon, S. Control of avian influenza infections in poultry with emphasis on vaccination. Expert Rev Anti Infect Ther 4, 751–757, doi:10.1586/14787210.4.5.751 (2006).

25 Steel, J. et al. Live attenuated influenza viruses containing NS1 truncations as vaccine candidates against H5N1 highly pathogenic avian influenza. J Virol 83, 1742–1753, doi:10.1128/JVI.01920-08 (2009).

26 Beug, H., von Kirchbach, A., Doderlein, G., Conscience, J. F. & Graf, T. Chicken hematopoietic cells transformed by seven strains of defective avian leukemia viruses display three distinct phenotypes of differentiation. Cell 18, 375–390, doi:10.1016/0092-8674(79)90057-6 (1979).

27 Amorim, M. J. et al. A Rab11- and microtubule-dependent mechanism for cytoplasmic transport of influenza A virus viral RNA. J Virol 85, 4143–4156, doi:10.1128/JVI.02606-10 (2011).

28 de Wit, E. et al. Efficient generation and growth of influenza virus A/PR/8/34 from eight cDNA fragments. Virus Res 103, 155–161, doi:10.1016/j.virusres.2004.02.028 (2004).

29 Jiang, H., Yu, K. & Kapczynski, D. R. Transcription factor regulation and cytokine expression following in vitro infection of primary chicken cell culture with low pathogenic avian influenza virus. Virol J 10, 342, doi:10.1186/1743-422X-10-342 (2013).

30 Spackman, E. & Killian, M. L. Avian influenza virus isolation, propagation, and titration in embryonated chicken eggs. Methods Mol Biol 1161, 125–140, doi:10.1007/978-1-4939-0758-8_12 (2014).

31 Noton, S. L. et al. Identification of the domains of the influenza A virus M1 matrix protein required for NP binding, oligomerization and incorporation into virions. J Gen Virol 88, 2280–2290, doi:10.1099/vir.0.82809-0 (2007).

32 Amorim, M. J., Read, E. K., Dalton, R. M., Medcalf, L. & Digard, P. Nuclear export of influenza A virus mRNAs requires ongoing RNA polymerase II activity. Traffic 8, 1–11, doi:10.1111/j.1600-0854.2006.00507.x (2007).

33 Schneider, C. A., Rasband, W. S. & Eliceiri, K. W. NIH Image to ImageJ: 25 years of image analysis. Nat Methods 9, 671–675, doi:10.1038/nmeth.2089 (2012).

34 Kapczynski, D. R. et al. Characterization of the 2012 highly pathogenic avian influenza H7N3 virus isolated from poultry in an outbreak in Mexico: pathobiology and vaccine protection. J Virol 87, 9086–9096, doi:10.1128/JVI.00666-13 (2013).

35 Bertran, K. et al. Protection of White Leghorn chickens by U.S. emergency H5 vaccination against clade 2.3.4.4 H5N2 high pathogenicity avian influenza virus. Vaccine 35, 6336-6344, doi:10.1016/j.vaccine.2017.05.051 (2017).

36 Spackman, E. et al. Development of real-time RT-PCR for the detection of avian influenza virus. Avian Dis 47, 1079–1082, doi:10.1637/0005-2086-47.s3.1079 (2003).

37 Pedersen, J. C. Hemagglutination-inhibition test for avian influenza virus subtype identification and the detection and quantitation of serum antibodies to the avian influenza virus. Methods Mol Biol 436, 53–66, doi:10.1007/978-1-59745-279-3_8 (2008).

38 Kumar, S., Stecher, G. & Tamura, K. MEGA7: Molecular Evolutionary Genetics Analysis Version 7.0 for Bigger Datasets. Mol Biol Evol 33, 1870–1874, doi:10.1093/molbev/msw054 (2016).

39 Deshpande, K. L., Fried, V. A., Ando, M. & Webster, R. G. Glycosylation affects cleavage of an H5N2 influenza virus hemagglutinin and regulates virulence. Proc Natl Acad Sci U S A 84, 36–40, doi:10.1073/pnas.84.1.36 (1987).

40 Webster, R. G., Kawaoka, Y. & Bean, W. J., Jr. Molecular changes in A/Chicken/Pennsylvania/83 (H5N2) influenza virus associated with acquisition of virulence. Virology 149, 165–173, doi:10.1016/0042-6822(86)90118-2 (1986).

41 Zhang, M., Zharikova, D., Mozdzanowska, K., Otvos, L. & Gerhard, W. Fine specificity and sequence of antibodies directed against the ectodomain of matrix protein 2 of influenza A virus. Mol Immunol 43, 2195–2206, doi:10.1016/j.molimm.2005.12.015 (2006).

42 Himly, M., Foster, D. N., Bottoli, I., Iacovoni, J. S. & Vogt, P. K. The DF-1 chicken fibroblast cell line: transformation induced by diverse oncogenes and cell death resulting from infection by avian leukosis viruses. Virology 248, 295–304, doi:10.1006/viro.1998.9290 (1998).

43 Roberts, P. C., Lamb, R. A. & Compans, R. W. The M1 and M2 proteins of influenza A virus are important determinants in filamentous particle formation. Virology 240, 127–137, doi:10.1006/viro.1997.8916 (1998).

44 Rossman, J. S. et al. Influenza virus m2 ion channel protein is necessary for filamentous virion formation. J Virol 84, 5078–5088, doi:10.1128/JVI.00119-10 (2010).

45 Beale, R. et al. A LC3-interacting motif in the influenza A virus M2 protein is required to subvert autophagy and maintain virion stability. Cell Host Microbe 15, 239–247, doi:10.1016/j.chom.2014.01.006 (2014).

46 Elton, D. et al. The genetics of virus particle shape in equine influenza A virus. Influenza Other Respir Viruses 7 **Suppl 4**, 81–89, doi:10.1111/irv.12197 (2013).

47 Campbell, P. J. et al. The M segment of the 2009 pandemic influenza virus confers increased neuraminidase activity, filamentous morphology, and efficient contact transmissibility to A/Puerto Rico/8/1934-based reassortant viruses. J Virol 88, 3802–3814, doi:10.1128/JVI.03607-13 (2014).

48 Seladi-Schulman, J., Steel, J. & Lowen, A. C. Spherical influenza viruses have a fitness advantage in embryonated eggs, while filament-producing strains are selected in vivo. J Virol 87, 13343–13353, doi:10.1128/JVI.02004-13 (2013).

49 Lakdawala, S. S. et al. Eurasian-origin gene segments contribute to the transmissibility, aerosol release, and morphology of the 2009 pandemic H1N1 influenza virus. PLoS Pathog 7, e1002443, doi:10.1371/journal.ppat.1002443 (2011).

50 Yi, Z. et al. Exploring the determinants of influenza A/H7N9 control intervention efficacy in China: Disentangling the effect of the ’1110’ policy and poultry vaccination. Transbound Emerg Dis 69, e1982–e1991, doi:10.1111/tbed.14532 (2022).

51 Swayne, D. E. & Kapczynski, D. Strategies and challenges for eliciting immunity against avian influenza virus in birds. Immunol Rev 225, 314–331, doi:10.1111/j.1600-065X.2008.00668.x (2008).

52 Sugrue, R. J. et al. Specific structural alteration of the influenza haemagglutinin by amantadine. EMBO J 9, 3469–3476, doi:10.1002/j.1460-2075.1990.tb07555.x (1990).

53 Ohuchi, M. et al. Rescue of vector-expressed fowl plague virus hemagglutinin in biologically active form by acidotropic agents and coexpressed M2 protein. J Virol 68, 920–926, doi:10.1128/JVI.68.2.920-926.1994 (1994).

54 Ciampor, F. et al. Evidence that the amantadine-induced, M2-mediated conversion of influenza A virus hemagglutinin to the low pH conformation occurs in an acidic trans Golgi compartment. Virology 188, 14-24, doi:10.1016/0042-6822(92)90730-d (1992).

55 Ciampor, F., Thompson, C. A., Grambas, S. & Hay, A. J. Regulation of pH by the M2 protein of influenza A viruses. Virus Res 22, 247–258, doi:10.1016/0168-1702(92)90056-f (1992).

56 Germeraad, E. A. et al. Virus Shedding of Avian Influenza in Poultry: A Systematic Review and Meta-Analysis. Viruses 11, doi:10.3390/v11090812 (2019).

57 Pantin-Jackwood, M. J. et al. Efficacy of a Recombinant Turkey Herpesvirus H5 Vaccine Against Challenge With H5N1 Clades 1.1.2 and 2.3.2.1 Highly Pathogenic Avian Influenza Viruses in Domestic Ducks (Anas platyrhynchos domesticus). Avian Dis 60, 22-32, doi:10.1637/11282-091615-Reg.1 (2016).

58 Seo, S. H., Peiris, M. & Webster, R. G. Protective cross-reactive cellular immunity to lethal A/Goose/Guangdong/1/96-like H5N1 influenza virus is correlated with the proportion of pulmonary CD8(+) T cells expressing gamma interferon. J Virol 76, 4886–4890, doi:10.1128/jvi.76.10.4886-4890.2002 (2002).

59 Seo, S. H. & Webster, R. G. Cross-reactive, cell-mediated immunity and protection of chickens from lethal H5N1 influenza virus infection in Hong Kong poultry markets. J Virol 75, 2516–2525, doi:10.1128/JVI.75.6.2516-2525.2001 (2001).

60 Bradshaw, G. L., Schwartz, C. D. & Schlesinger, R. W. Replication of H1N1 influenza viruses in cultured mouse embryo brain cells: virus strain and cell differentiation affect synthesis of proteins encoded in RNA segments 7 and 8 and efficiency of mRNA splicing. Virology 176, 390–402, doi:10.1016/0042-6822(90)90009-g (1990).

